# Attenuated Subcomponent Vaccine Design Targeting the SARS-CoV-2 Nucleocapsid Phosphoprotein RNA Binding Domain: *In silico* analysis

**DOI:** 10.1101/2020.06.30.176537

**Authors:** Onyeka S. Chukwudozie, Rebecca C. Chukwuanukwu, Iroanya O. Onyekachi, Eze M. Daniel, Duru C. Vincent, Dele-Alimi T. Onaopemipo, Busuyi D. Kehinde, Bankole T. Taiwo, Obi C. Perpetua, Okinedo U. Elizabeth

**Affiliations:** Department of Cell Biology and Genetics, University of Lagos, Akoka Lagos state, Nigeria; Immunology Unit, Medical Laboratory Science Department, Nnamdi Azikiwe University, Nnewi Campus; Molecular Genetics unit, Institute of Child Health, College of Medicine, University of Ibadan, Oyo state, Nigeria; Department of Biochemistry, Ladoke Akintola University of Technology, Ogbomosho, Oyo State, Nigeria; Immunology and Bioinformatics unit, Department of Parasitology and Entomology, Nnamdi Azikiwe University, Awka, Anambra state, Nigeria; Department of Science Laboratory and Technology (Microbiology Unit), Federal Polytechnic, Oko, Anambra State, Nigeria

**Keywords:** COVID-19, Epitope Vaccine, Reverse vaccinology, Molecular dynamics simulation, Nucleocapsid protein

## Abstract

The novel coronavirus disease (COVID-19) caused by severe acute respiratory syndrome coronavirus 2 (SARS-CoV-2) has previously never been identified with humans, thereby creating devastation in public health. The need for an effective vaccine to curb this pandemic cannot be overemphasized. In view of this, we, therefore, designed a subcomponent antigenic peptide vaccine targeting the N-terminal (NT) and C-terminal (CT) RNA binding domains of nucleocapsid protein that aid in viral replication. Promising antigenic B-cells and T cell epitopes were predicted using computational pipelines. The peptides “RIRGGDGKMKDL” and “AFGRRGPEQTQGNFG” were the B cell linear epitopes with good antigenic index and non-allergenic property. Two CD8^+^ and Three CD4^+^ T-cell epitopes were also selected considering their safe immunogenic profiling such as allergenicity, antigen level conservancy, antigenicity, peptide toxicity, and putative restrictions to a number of MHC-I and II alleles. With these selected epitopes, a non-allergenic chimeric peptide vaccine incapable of inducing a Type II hypersensitivity reaction was constructed. The molecular interaction between the toll-like receptor-5 (TLR5) which was triggered by the vaccine was analyzed by molecular docking and scrutinized using dynamics simulation. Finally, *in silico* cloning was performed to ensure the expression and translation efficiency of the vaccine, utilizing pET-28a vector. This research, therefore, provides a guide for experimental investigation and validation.

## Introduction

Coronavirus disease 2019 (COVID-19) is the disease associated with severe acute respiratory syndrome coronavirus-2 (SARS-CoV-2). Characterizing COVID-19 as a pandemic is an acknowledgment that the coronavirus disease 2019 (COVID-19) outbreak, which started in the Hubei province of China in 2019, has now spread to all continents, affecting most countries around the world with differential impacts and peculiarities [**1, 2**].

Coronaviruses have the largest known genomes (up to 32 kb) among +RNA viruses, and they encode four structural and sixteen non-structural proteins [**3**]. The non-structural proteins (nsp) consist of all the enzymatic activities that are imperative for the viral replication, mostly associated with RNA replication [**3, 4**]. Structurally, the SARS-CoV-2 genome also encodes an RNA-dependent RNA-polymerase complex consisting of the nsp7, nsp8 and nsp12, the RNA capping machinery which also constitute the nsp10, nsp13, nsp14 and nsp16, and finally additional enzymes such as proteases (the nsp3 PLpro and the nsp5 3CLpro) which cleave viral polyproteins and/or impede innate immunity [**5, 6, 7**].

The four structural proteins together with the viral +RNA genome and the envelope constitute the virion [**5, 8**]. The matrix (M), small envelope (E), and spike (S) proteins are embedded within the lipid envelope [**6, 9**]. The fourth structural protein, the nucleocapsid phosphoprotein (N), physically links the envelope to the +RNA genome. It consists of an N-terminal (NTD) and a C-terminal (CTD) domain [**10, 11**]. Both domains are capable of RNA binding. In addition, the CTD serves as a dimerization domain and binds the matrix protein forming the physical link between the+ RNA genome and the envelope [**12**]. The SARS N protein has also been shown to modulate the host intracellular machinery and play regulatory roles during the viral life cycle [**5-7**]. In light of the genomic similarities between SARS-CoV and SARS-CoV-2, it is reasonable to expect the N protein to function in a similar way [**10**]. All the SARS-CoV-2 proteins are potential drug and vaccine targets and a detailed understanding of their functions is therefore of utmost importance. The nucleocapsid phosphoprotein of the severe acute respiratory syndrome coronavirus N (SARS-CoV N) protein packages the viral genome into a helical ribonucleocapsid (RNP) and plays a fundamental role during viral self-assembly [**5-7**]. It is a protein with multifarious activities. Furthermore, N protein is frequently used in vaccine development and serological assays [**13**]. At present, few reports focus on SARS-CoV-2 N protein, and there is an urgent need for an updated understanding of SARS-CoV-2 N protein. Majorly, the vaccine therapeutic experiments are highly centered on the spike or entire protein, but we are focusing mainly on the nucleocapsid phosphoprotein which is a protein subset of the virus.

## Materials and Methods

### Data retrieval, Structural and Physiochemical Analysis of SARS-CoV-2 Nucleocapsid Protein

The protein sequence of the SARS-CoV-2 nucleocapsid phosphoprotein was retrieved in a FASTA format from the NCBI repository with the accession numbers: Wuhan, China (Genbank ID: QHD43423.2). The X-ray crystal structure of the nucleocapsid protein was also retrieved from the protein data bank (PDB: 6m3m). Retrieved structure was subjected to a structural alignment to ascertain the level of homology and probable mutations that have occurred over time during viral replication among the coronavirus family. Bootstrap value and other default parameters were used to fabricate the alignment. The physiochemical properties of the protein sequence were assessed and bio-computed via an online tool Protparam **[14]** (http://web.expasy.org/protparam/).

### Putative B Cell Linear and Discontinuous Epitopes

The Nucleocapsid sequence was analyzed with a view to recognize the antigenic regions that were achieved by predicting epitopic peptides. The promising antigenic linear B cell epitopes were predicted using the Bepipred server from the Immune-Epitope-Database and Analysis-Resource (IEDB) database [**15**]. BepiPred-2.0 is based on a random forest algorithm trained on epitopes annotated from antibody-antigen protein structures [**15**]. About 12-15 mers (residues) was assumed to bind to MHC groove. Other criteria such as antigenicity, surface accessibility, flexibility, and hydrophobicity were considered as part of the profiling process of the antigenic B cell linear epitopes. For antigenicity testing, these epitopes were subjected to Vaxijen 2.0 server at a threshold of 0.6 **[16]**. The next stage of screening was the prediction of the discontinuous epitopes which are folded in conformation aiding in the antibody recognition of denatured antigens. The Elliprot server (http://tools.iedb.org/ellipro/) was adopted for this purpose, while Pymol was utilized to examine the positions of forecast epitopes on the 3D structure of SARS-CoV-2 nucleocapsid phosphoprotein [**17**].

### Prediction of Cytotoxic T lymphocytes (CTL) and Helper T lymphocytes (HTL) epitopes

The CTL epitopes were predicted using the IEDB MHC I binding prediction algorithms (http://tools.iedb.org/mhci). This method integrates the prediction of epitopes restricted to a large number of MHC I alleles and proteasomal C-terminal cleavage, using artificial neural network application. For better predictive accuracy, other software such as artificial neural network (ANN), stabilized matrix method (SMM), MHC binding energy covariance matrix (SMMPMBEC), NetMHCpan, pickpocket, and NetMHCstapan, were adopted for this purpose. To predict the HTL cell epitopes, the MHC II binding predictions tool (http://tools.iedb.org/mhcii/) found in the IEDB database was adopted. The antigenic properties of the epitopes were studied using the Vaxijen 2.0 server set at a threshold of 0.6. The peptide toxicity predicted from the ToxinPred server (http://crdd.osdd.net/raghava/toxinpred/), allergenicity, predicted from AllergenFP 1.0 and digestion predicted from protein digest server, were all considered in selecting the final epitopes.

### Prediction of the 3D Structures of the Predicted Epitopes and HLA-A 0201 Allele for Molecular Docking

The molecular docking of the antigenic epitopic peptides was conducted with the alleles they were mostly restricted to, which was HLA-A 0201. The protein structure of the allele was retrieved from the protein data bank with the identifier (PDB: 4U6Y), while the predicted peptides 3D structures were modeled via PEPFOLD server at RPBS MOBYL portal. The best models provided by the server were chosen for the docking study. The HAWKDOCK server was employed for the docking process. It combines ATTRACT for global macromolecular docking and HawKRank for scoring.

### Homology modeling of the conjugated peptide vaccine

The three-dimensional model of the conjugated antigenic vaccine was predicted using the I-TASSER server which generates a 3D model of query sequence by multiple threading alignments and iterative structural assembly simulation [**18].** The I-TASSER online server was selected for its availability, composite approach of modeling, and performance in CASP competition. The quality of generated 3D models was checked by Z-score and the best model is selected for further consideration. The functional analogs were ranked based on TM-score, RMSD, sequence identity, and coverage of the structure alignment. The quality of the predicted model was determined by C-score (confidence score) which is ranged as −5 to 2. The obtained 3D model of the conjugated antigenic vaccine and the human Toll-like receptor-5 PDB structures were aligned employing the TM-align [**19**]. A quick and accurate structural alignment tool for two protein structures of unknown equivalence. An optimal superposition of the two structures built on the detected alignment, as well as the TM-score value which scales the structural similarity. TM-score has the value in (0,1), where 1 indicates a perfect match between two structures.

### Validation of predicted conjugated Peptide Vaccine 3D model

The confirmation of the selected 3D model predicted by I-TASSER was further validated by the Ramachandran plot. RAMPAGE and MolProbity [**20**] online servers were employed for the estimation of selected 3D model quality. It can begin from either C-alpha trace, main-chain model, or full-atomic model. The Ramachandran plot obtained from RAMPAGE describes a good quality model that has over 70% residues in the most favored region. The plot analysis was able to show the allowed and disallowed dihedral angles psi (ψ) and phi (ϕ) of an amino acid which is calculated based on van der Waal radius of the side chain. The corresponding percentage value of both the allowed and disallowed region of the separate plots of glycine and proline residues of the modeled structure was generated. Qualitative evaluation of 3D models was employed by ProSA [**21**]. ProSA specifically faces the needs confronted in the authentication of protein structures acquired from X-ray analysis, NMR spectroscopy, and hypothetical estimations.

### Protein-protein docking of the peptide vaccine and the human toll-like receptor-5 (TLR5)

In this study, molecular docking analysis between vaccine and human toll-like receptor-5 was performed using ClusPro 2.2 protein-protein interaction online server [**22**]. The shape complementarity and minimal binding energy of Toll-like receptor-5 with predicted conjugated antigenic vaccine model obtained from I-TASSER, is determined by the cluster scores for lowest binding energy prediction, calculated using the formula [**23**]:

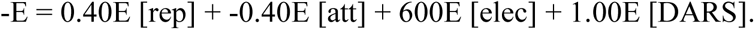

Here, repulsive, attractive, electrostatic as well as interactions extracted from the decoys as the reference state, are considered for structure-based pairwise potential calculation in docking [**22**]. The best PDB conformation was subjected to Prodigy server to ascertain the binding free energy of the protein complex.

### Molecular Dynamics Simulations

The interacting complex between the vaccine and the toll-like receptor (PDB: 3J0A), was thoroughly accessed based on the existing coordinates between the docked protein complex. Parameters considered were the deformability, B factor, eigenvalues associated with the normal mode which represents the motion stiffness. The lower the eigenvalue, the easier the deformation. The covariance matrix was also considered for the simulation. It indicates the coupling between the pairs of residues. The correlation matrix is computed using the Cα Cartesian coordinates **[24]**.

### Codon Optimization and *In silico* Cloning

A codon optimization was conducted to ascertain the maximum expression of the vaccine in the host. This was done with the aim of boosting the vaccine translational rate in *E. coli* K12. Restriction enzymes cleavage sites, prokaryote ribosomal binding site, and finally rho-independent transcription termination, were all avoided during the option selection. Codon adaptation index (CAI) value and GC content of the adapted sequence was obtained and compared with the ideal range. The obtained refined nucleotide was cloned into the pET28a D315A vector.

## Results

### Structural alignment Studies of the SARS-CoV-2 NP

The protein structure consists of 4 side chains as shown, which suggests a plausible model for RNA binding [**Figure 1**]. A structural alignment was performed to ascertain the level of conservancy across the coronavirus NP, while the SARS-CoV-2 (PDB: 6m3m) was maintained as the reference protein. Sequence similarity search and multiple sequence alignment (MSA) was adopted for this purpose. The MSA of the 14 homologous proteins to 6m3m was generated with a BLAST search against the PDBAA database. Across the sequence alignment, the various mutations including deletions supersede the conserved regions of the amino residues [**Figure 2**]. In summary, the alignment construct was less conserved, signifying the rapid rate of mutations that has acted on the protein during viral replications. The protein identification of the PDB ID’s are summarized [**supplementary data**]. Also, results of the solvent accessibility and hydropathy of the N NTD of coronaviruses shows that most of the regions on the sequence have high solvent accessibility and are hydrophilic, properties which makes the N NTD a likely antigenic target.

**Figure 1:**
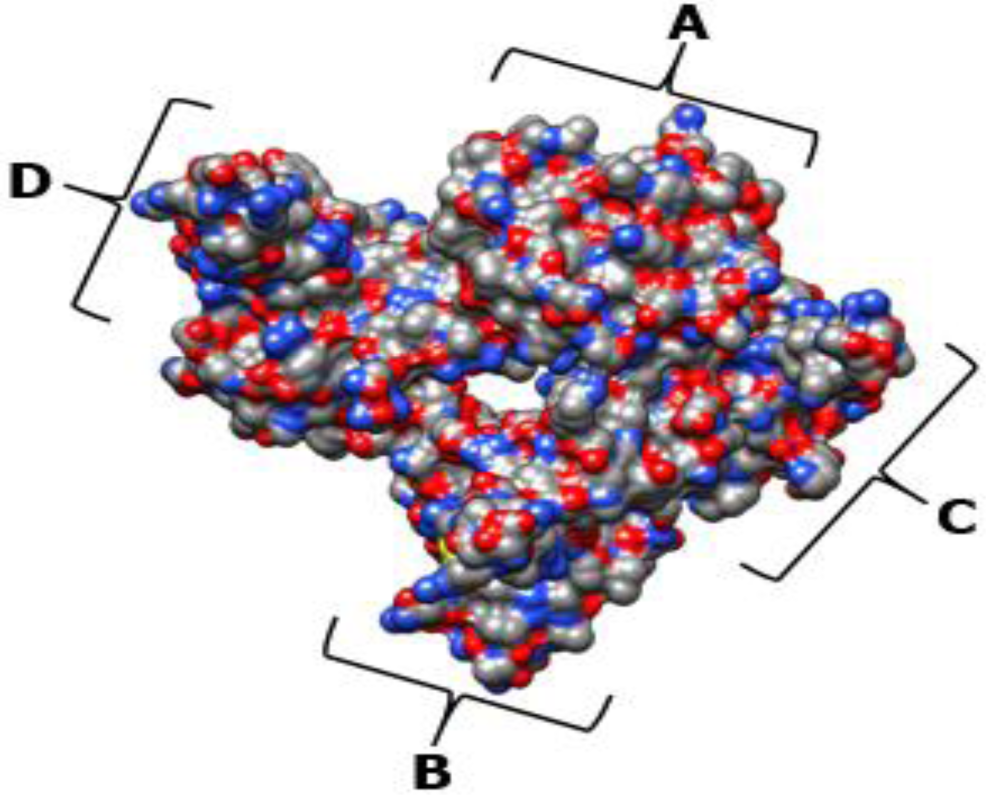
Crystal structure of SARS-CoV-2 nucleocapsid protein N-terminal RNA binding domain at a resolution of 2.70 Å (PDB: 6m3m). The chain components are identified respectively.

**Figure 2:**
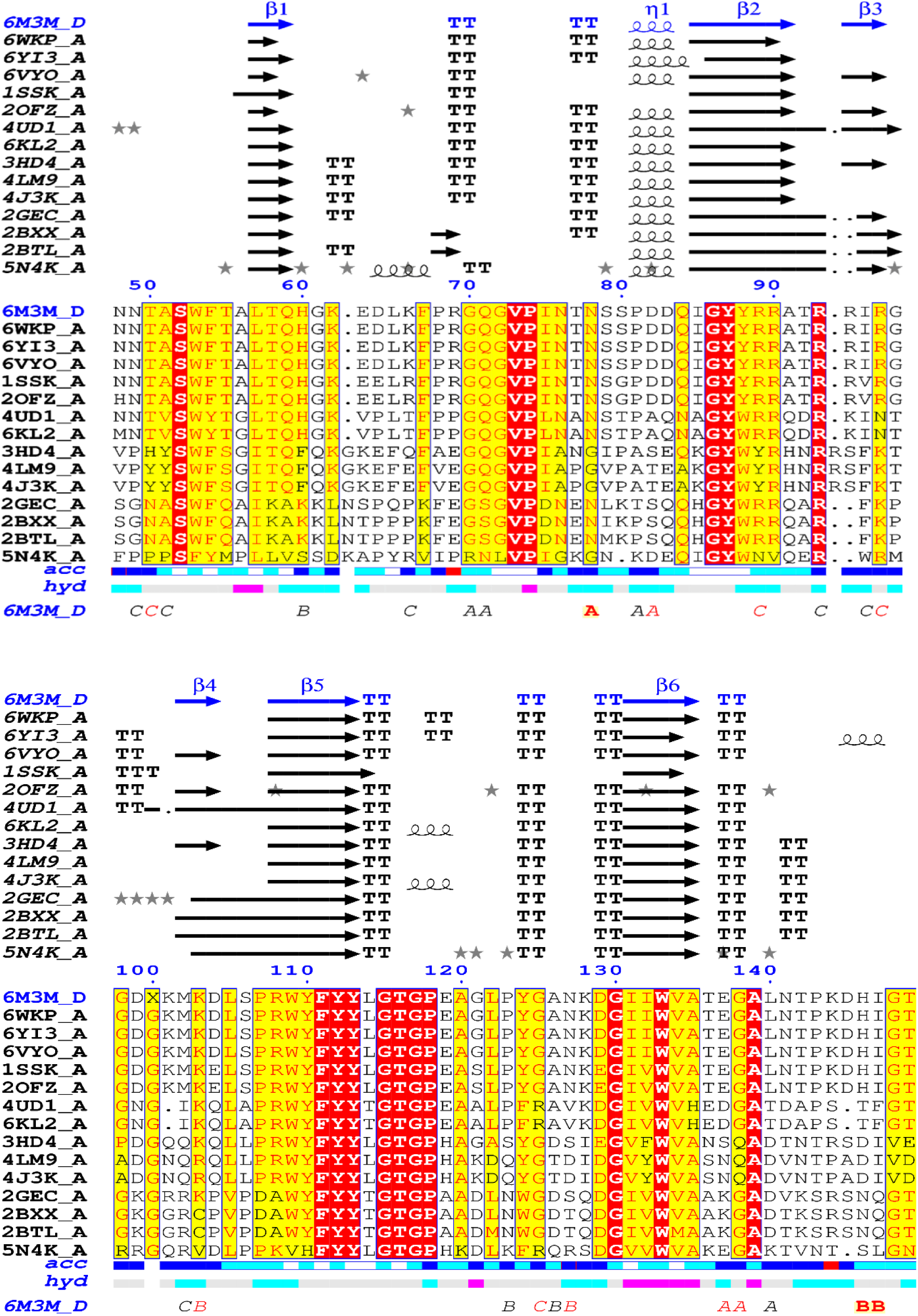
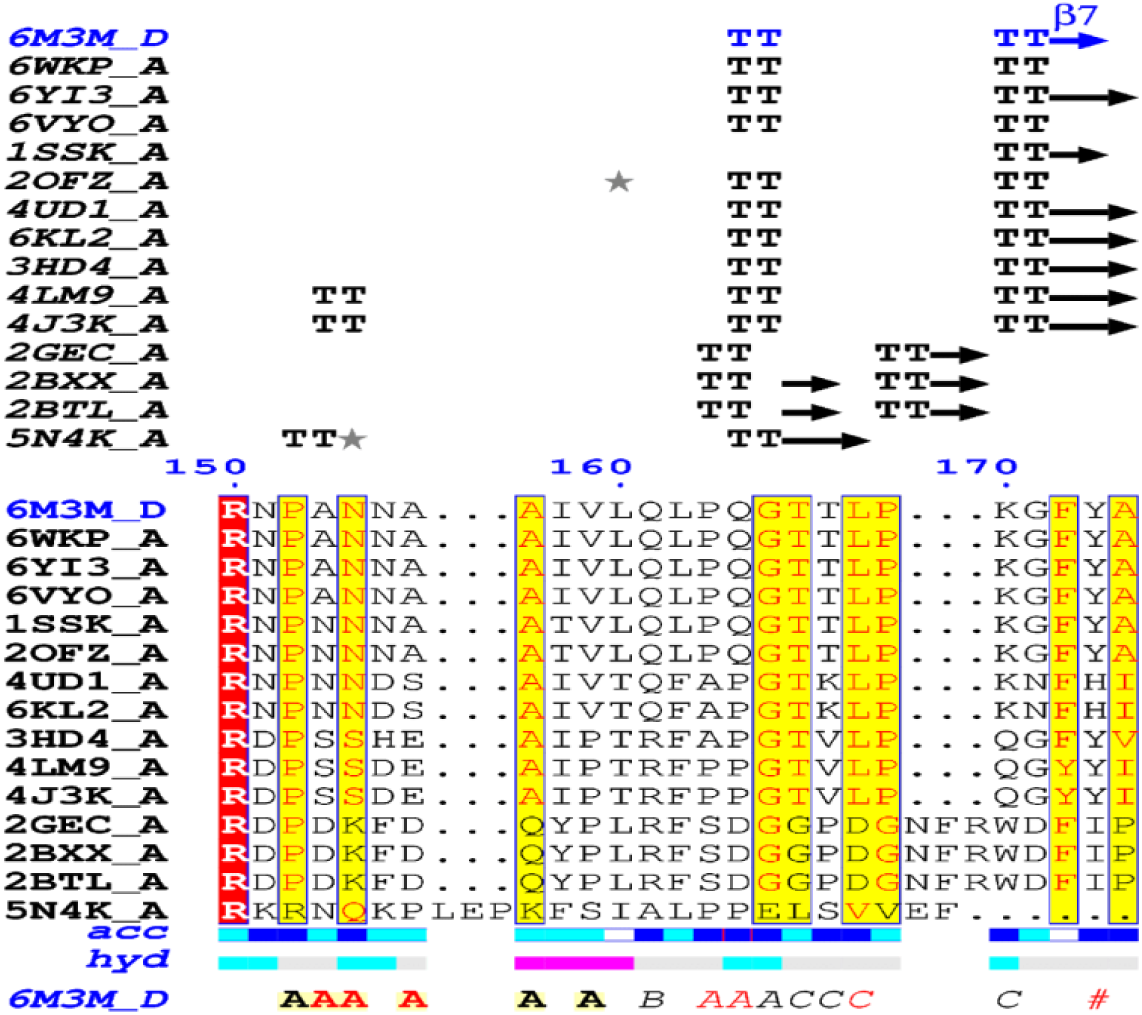
Structural alignment of the SARS-CoV-2 RNA binding domain of nucleocapsid phosphoprotein. Helices are represented in squiggles, while beta strands with arrows and turns with TT letters. Solvent accessibility is rendered by a first bar below the sequence (blue is accessible, cyan is intermediate, white is buried) and hydropathy by a second bar below (pink is hydrophobic, white is neutral, cyan is hydrophilic). Bottom letters and symbols depict crystallographic. Alignments in reds represent conserved regions, yellow highlights the regions that tend towards monomorphism, while white is regions that are highly mutated. The dotted segments are the sequence deletions.

### B cell linear and Discontinuous epitopes

Utilizing the Kolaskar and Tongaonkar antigenicity scale, Emini surface accessibility and the Chou and Fasman beta turn predictions, regions with viable antigenic properties were predicted. This scale was able to show the favorable regions across the protein that are potentially antigenic [**Figure 3a-3e**]. A total of four antigenic B cell epitopes was predicted. These epitopes had a safe physiochemical property such as the absence of peptide toxicity and lesser allergenicity, making it safe for vaccine production. Three of the selected epitopes had 100% across the antigen. Based on the conservancy across the antigen and the allergenicity, only RIRGGDGKMKDL and AFGRRGPEQTQGNFG were selected as the final promising antigenic B cell linear epitopes [**Table 1**]. These epitopes were mapped out from the protein structure [**Figure 4a**]. Based on the SARS-CoV-2 nucleocapsid protein (PDB: 6m3m), the discontinuous epitopes were predicted considering their propensity scores. The identified denatured antigens by the neutralizing antibody are highlighted **[Figure 4b-f]**. The residues are juxtaposed enabling the antibody to recognize the 3D dimensional structure. The chain D component of the protein had the highest denatured antigens with a total of 81 residues, followed by chain B, C, and A respectively in that hierarchical order as the number of residues and their respective rank scores are ranked **[Table 2]**. A higher propensity score of 0.745 means that only 25% of the D-chain residues are non-epitope residues predicted as part of the epitope. Also, the result of the linear B-cell epitope prediction correlated with the discontinuous epitope as the dodecapeptide epitope predicted in the B-cell linear epitope was also found to be in the D-chain component of the protein, further indicating the antigenicity of the D-chain monomer of the N NTD.

**Table 1:**
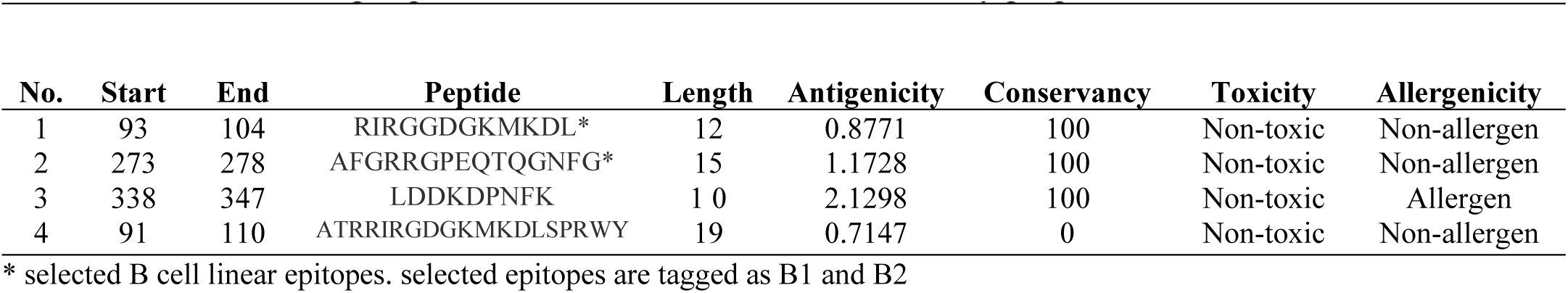
B cells linear epitopes of SARS-CoV-2 NP and their toxicity properties.

**Table 2:** The B cell discontinuous epitopes of the SARS-CoV-2 Nucleocapsid phosphoprotein

| No. | Residues | Number of residues | Score | 3d structure |
| --- | --- | --- | --- | --- |
| 1 | D:A56, D:L57, D:T58, D:Q59, D:H60, D:G61, D:K62, D:E63, D:D64, D:L65, D:K66, D:F67, D:P68, D:R69, D:G70, D:Q71, D:G72, D:V73, D:P74, D:Q84, D:Y88, D:R90, D:A91, D:T92, D:R93, D:R94, D:I95, D:R96, D:G97, D:G98, D:D99, D:K101, D:M102, D:K103, D:D104, D:L105, D:S106, D:P107, D:R108, D:W109, D:G117, D:P118, D:E119, D:A120, D:G121, D:L122, D:P123, D:Y124, D:G125, D:A126, D:N127, D:K128, D:D129, D:G130, D:I131, D:I132, D:W133, D:V134, D:A135, D:T136, D:E137, D:G138, D:A139, D:L140, D:N141, D:T142, D:P143, D:Q161, D:L162, D:P163, D:Q164, D:G165, D:T166, D:T167, D:L168, D:P169, D:K170, D:G171, D:F172, D:Y173, D:A174 | 81 | 0.745 | Figure 4b |
| 2 | B:N48, B:N49, B:T50, B:A51, B:S52, B:W53, B:F54, B:T55, B:A56, B:T58, B:Q59, B:H60, B:G61, B:P74, B:A91, B:T92, B:R93, B:R94, B:I95, B:R96, B:G97, B:D99, B:G100, B:K101, B:M102, B:K103, B:D104, B:L105, B:S106, B:P107, B:R108, B:Y110, B:L114, B:G115, B:T116, B:G117, B:P118, B:E119, B:A120, B:T142, B:P143, B:K144, B:D145, B:H146, B:I147, B:G148, B:T149, B:R150, B:N151, B:P152, B:A153, B:N154, B:N155, B:A156, B:A157, B:I158, B:V159, B:L160, B:G171 | 59 | 0.692 | Figure 4c |
| 3 | C:P47, C:N48, C:N49, C:T50, C:A51, C:W53, C:Q59, C:H60, C:G61, C:K62, C:E63, C:D64, C:L65, C:K66, C:F67, C:P68, C:R69, C:G70, C:Q71, C:G72, C:V73, C:S79, C:S80, C:P81, C:D82, C:D83, C:Q84, C:I85, C:G86, C:Y87, C:R89, C:R90, C:A91, C:T92, C:R93, C:R94, C:I95, C:R96, C:G98, C:D99, C:G100, C:K101, C:M102, C:K103, C:D104, C:L105, C:S106, C:P107, C:Y113, C:L114, C:G115, C:T116, C:G117, C:P118, C:E119, C:A120, C:G121, C:L122, C:P123, C:Y124, C:G125, C:A126, C:N127, C:K128, C:D129, C:G130, C:I131, C:I132, C:W133, C:V134, C:A135, C:T136, C:E137, C:G138, C:A139, C:L140, C:N141, C:T142, C:P143, C:K144, C:D145, C:H146, C:I147, C:G148, C:T149, C:R150, C:N151, C:P152, C:A153, C:N154, C:N155, C:T166, C:G171 | 93 | 0.683 | Figure 4d |
| 4 | A:H60, A:G61, A:K62, A:E63, A:D64, A:L65, A:K66, A:F67, A:P68, A:R69, A:G70, A:Q71, A:G72, A:V73, A:S79, A:S80, A:P81, A:D82, A:Q84, A:A135, A:T136, A:E137, A:G138, A:A139, A:L140, A:N141, A:T142, A:Q161, A:L162, A:P163, A:Q164, A:G165, A:T166, A:T167, A:L168, A:P169, A:K170, A:G171, A:Y173 | 39 | 0.658 | Figure 4e |
| 5 | B:I75, B:N76, B:T77, B:N78, B:S79, B:S80 | 6 | 0.617 | Figure 4f |

**Figure 3b-f:**
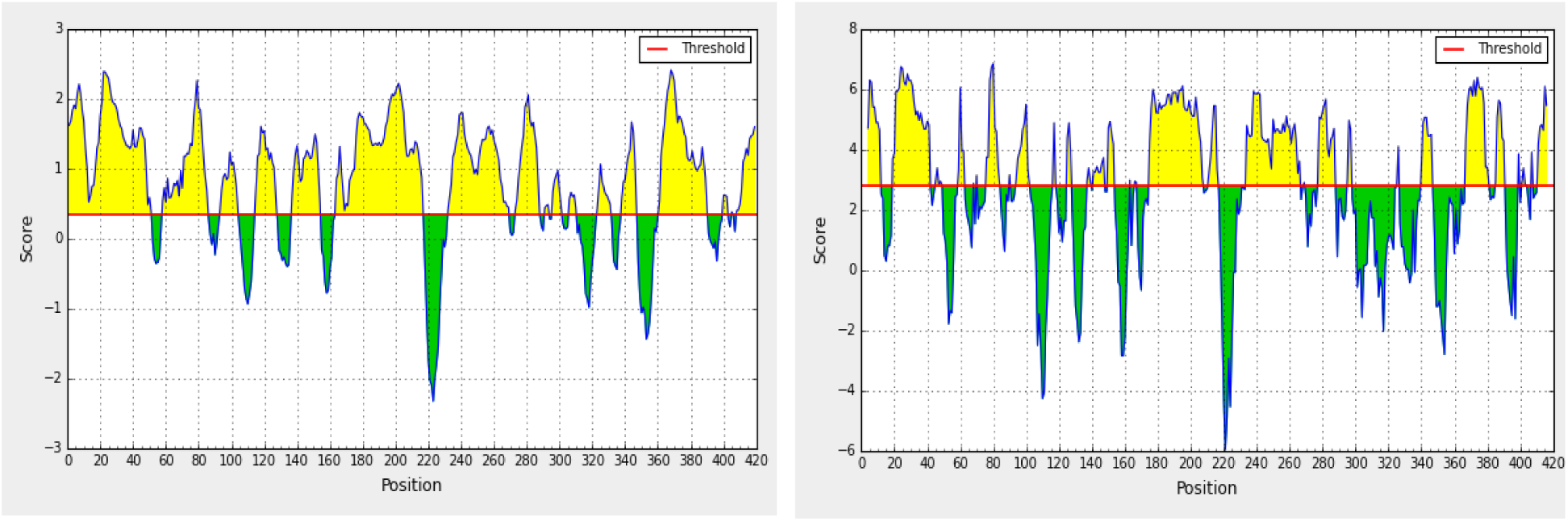
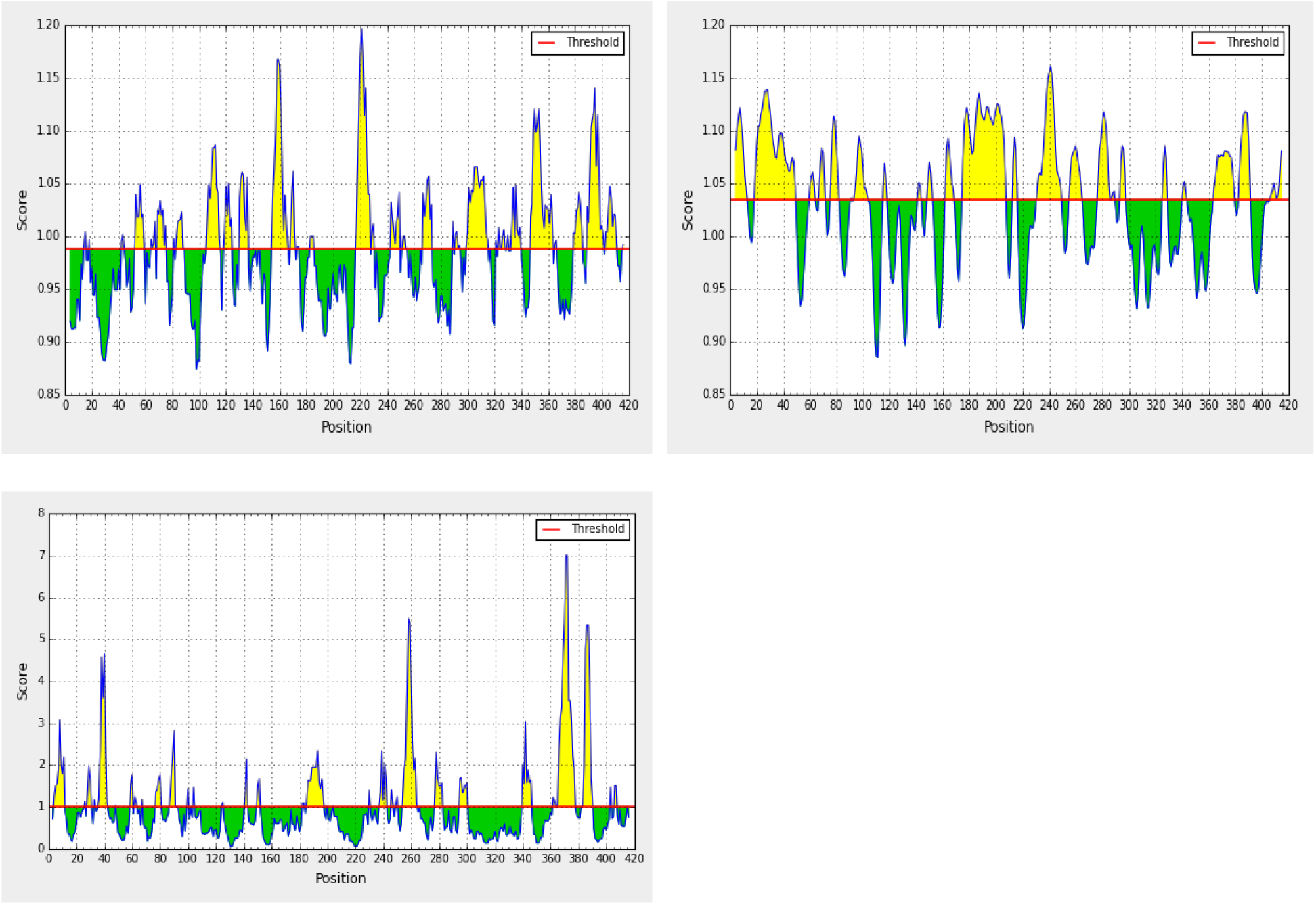
The properties of the B cell linear epitopes predictions. **a.** The Linear epitopes. **b.** hydrophobicity **c.** antigenicity **d.** flexibility and **e.** Surface accessibility. Green regions under the threshold color denote unfavorable related to the properties of interest. Yellow colors are above the threshold sharing higher scores. Horizontal red lines represent the threshold.

**Figure 4a:**
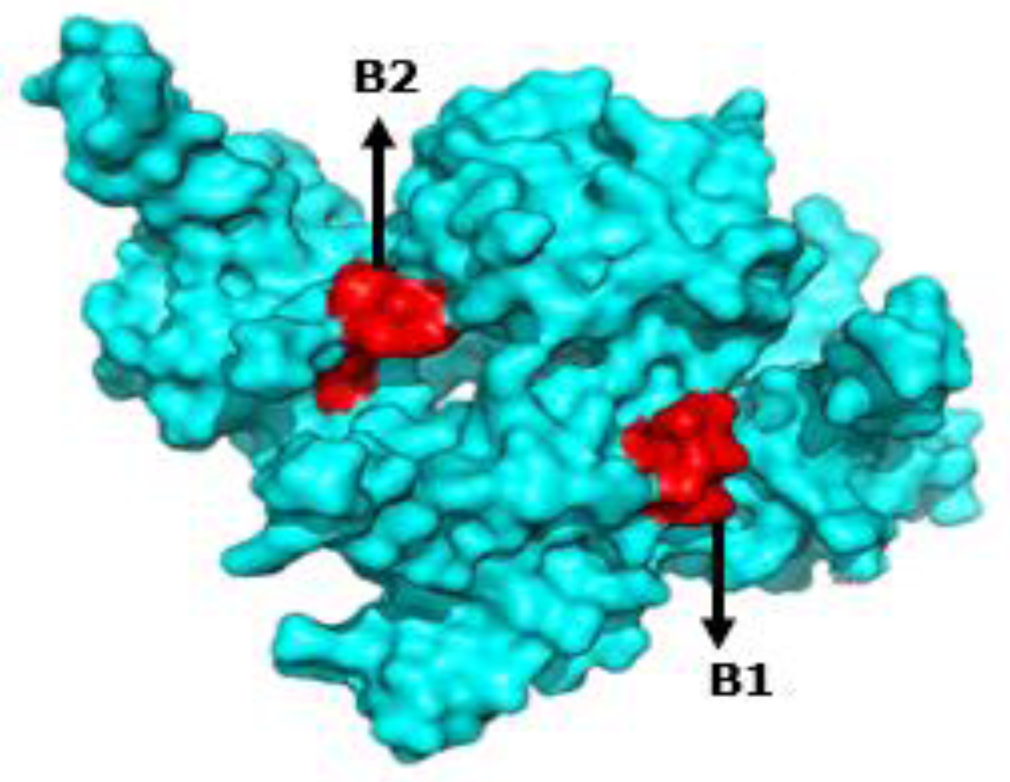
Mapped out antigenic B-cell linear epitopes (red) of the SARS-CoV-2 nucleocapsid protein

**Figure 4b-f:**
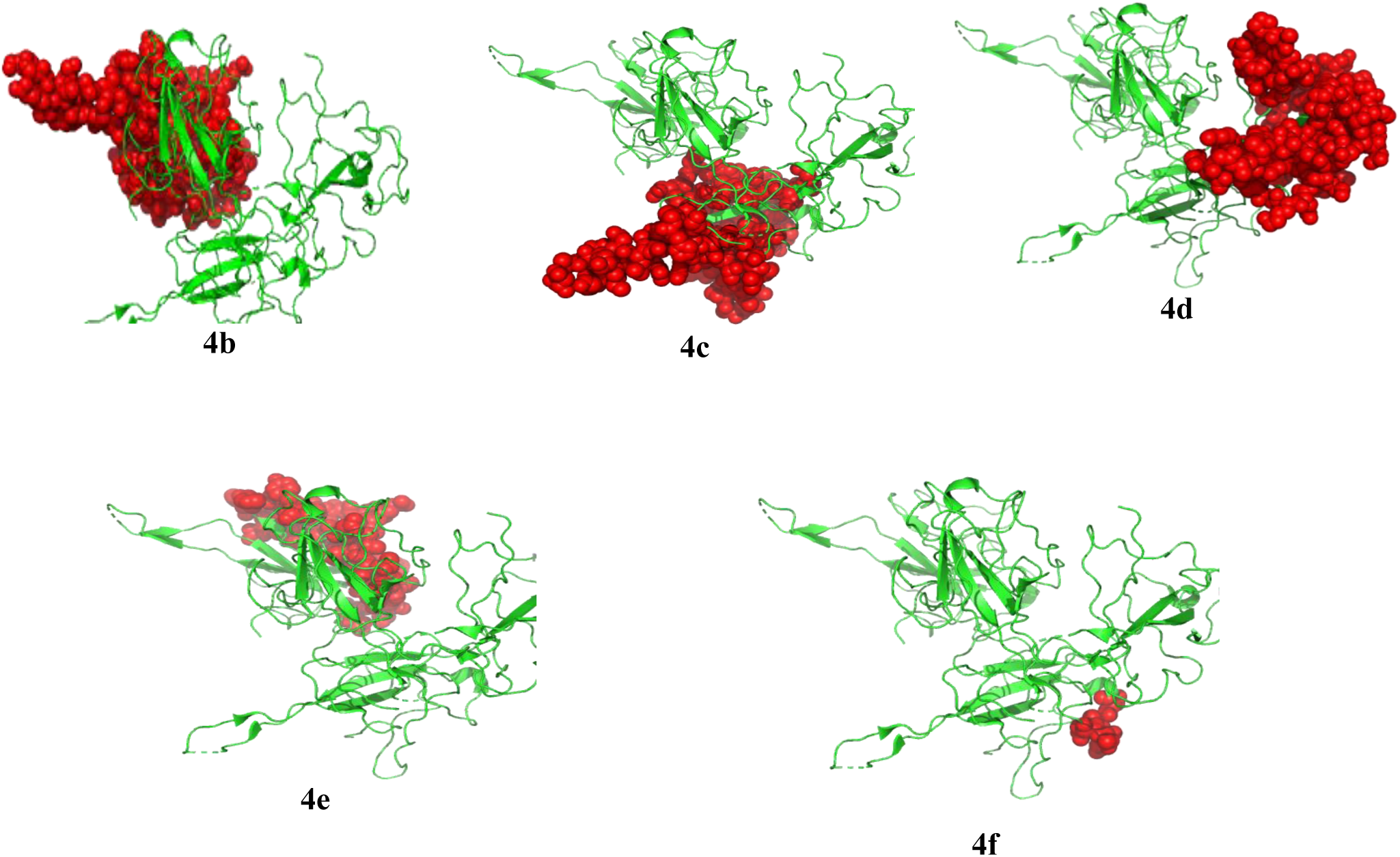
The antibody recognition of SARS-CoV-2 denatured antigens (red spheres) in **b**: D-monomer **c**: B Monomer **d**: C monomer **e**: A monomer **f**: B monomer respectively.

### The CTL and HTL Epitopes

Two non-allergenic and non-toxic cytotoxic epitopes, with a viable antigenic property, were selected. The epitopes are GMSRIGMEV and LTYTGAIKL [**Table 3**]. For the helper T cell epitopes, three were selected based on their conservancy score of 100%, non-allergenic attributes, and their respective antigenic properties [**Table 4**]. Few of the peptides, regardless of their antigenic nature, had an allergenic property capable of inducing a harmful autoimmune response. The promiscuity of the chosen peptides was also evaluated by considering the number of MHC-I and II alleles they are putatively restricted to.

**Table 3:**
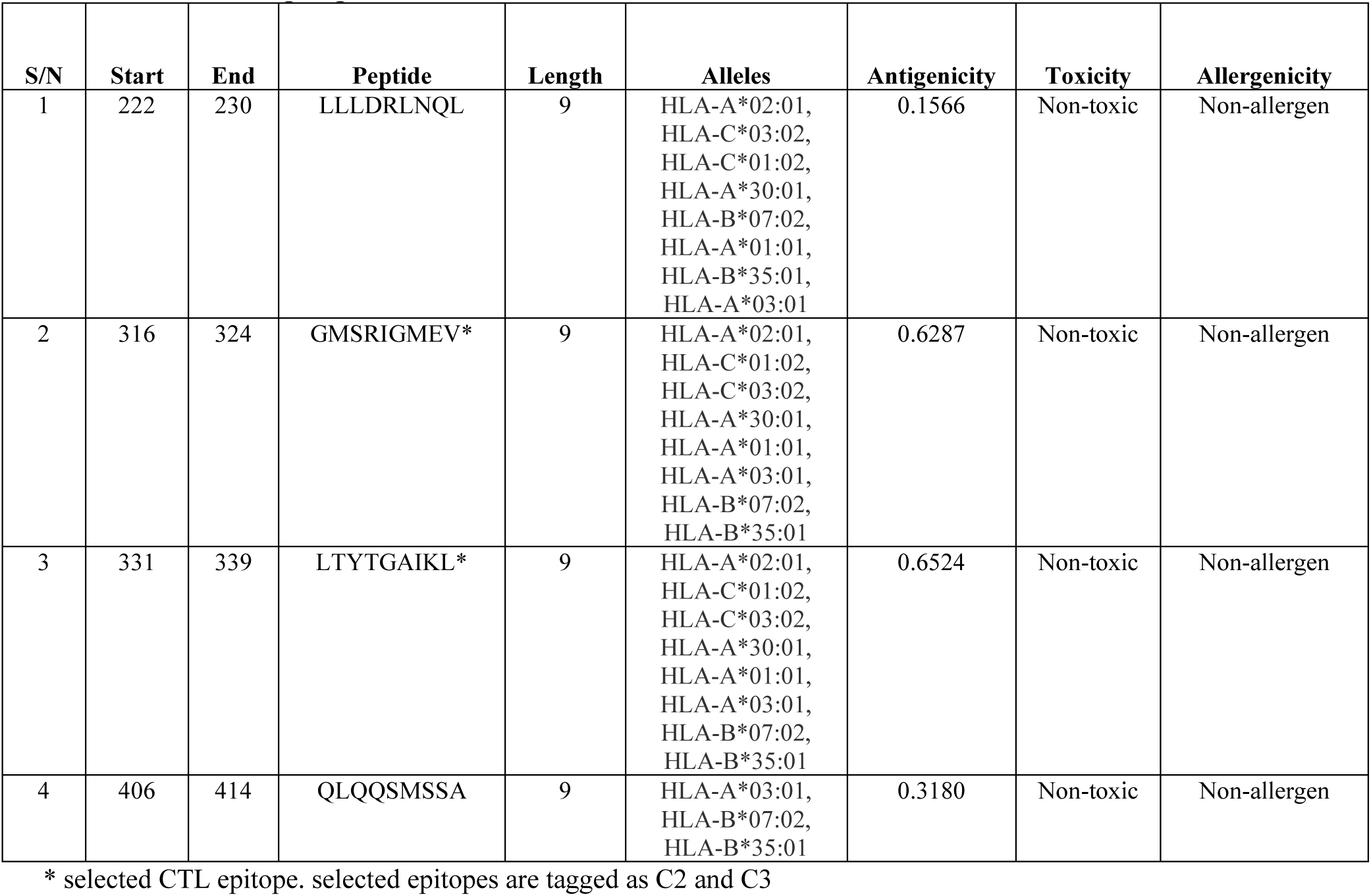
MHC-I epitopes of SARS-CoV-2 NP

**Table 4:**
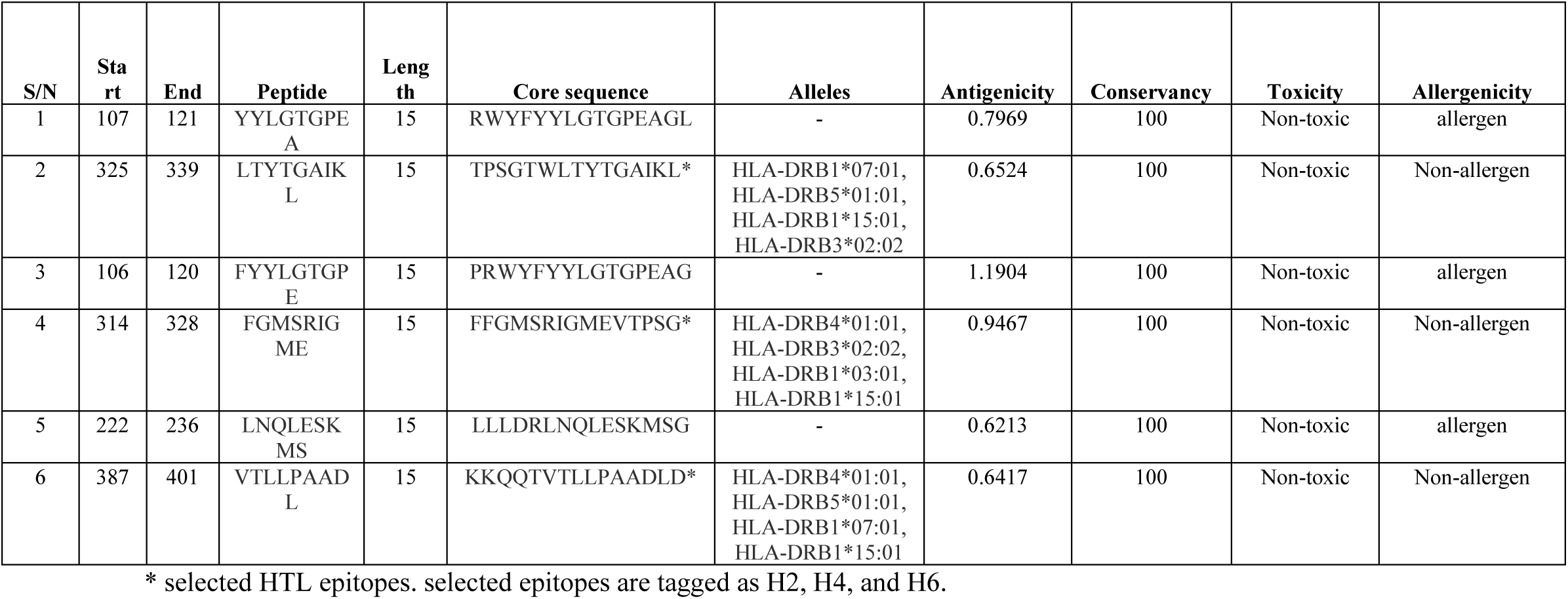
MHC-II epitopes of SARS-CoV-2 NP

### Molecular Docking of HLA-Epitope Interaction with the MHC-I Molecule

Both peptides “GMSRIGMEV and LTYTGAIKL” bind respectively to their restricted HLA molecules. Both epitopic peptides were highly restricted to several MHC-I molecules. In the case of HLA-A 0201, the binding free energy of GMSRIGMEV with the MHC-I antigen-binding groove was −8.3 kcal/mol, while the peptide LTYTGAIKL had binding energy of −10 kcal/mol respectively [**Figure 5**].

**Figure 5:**
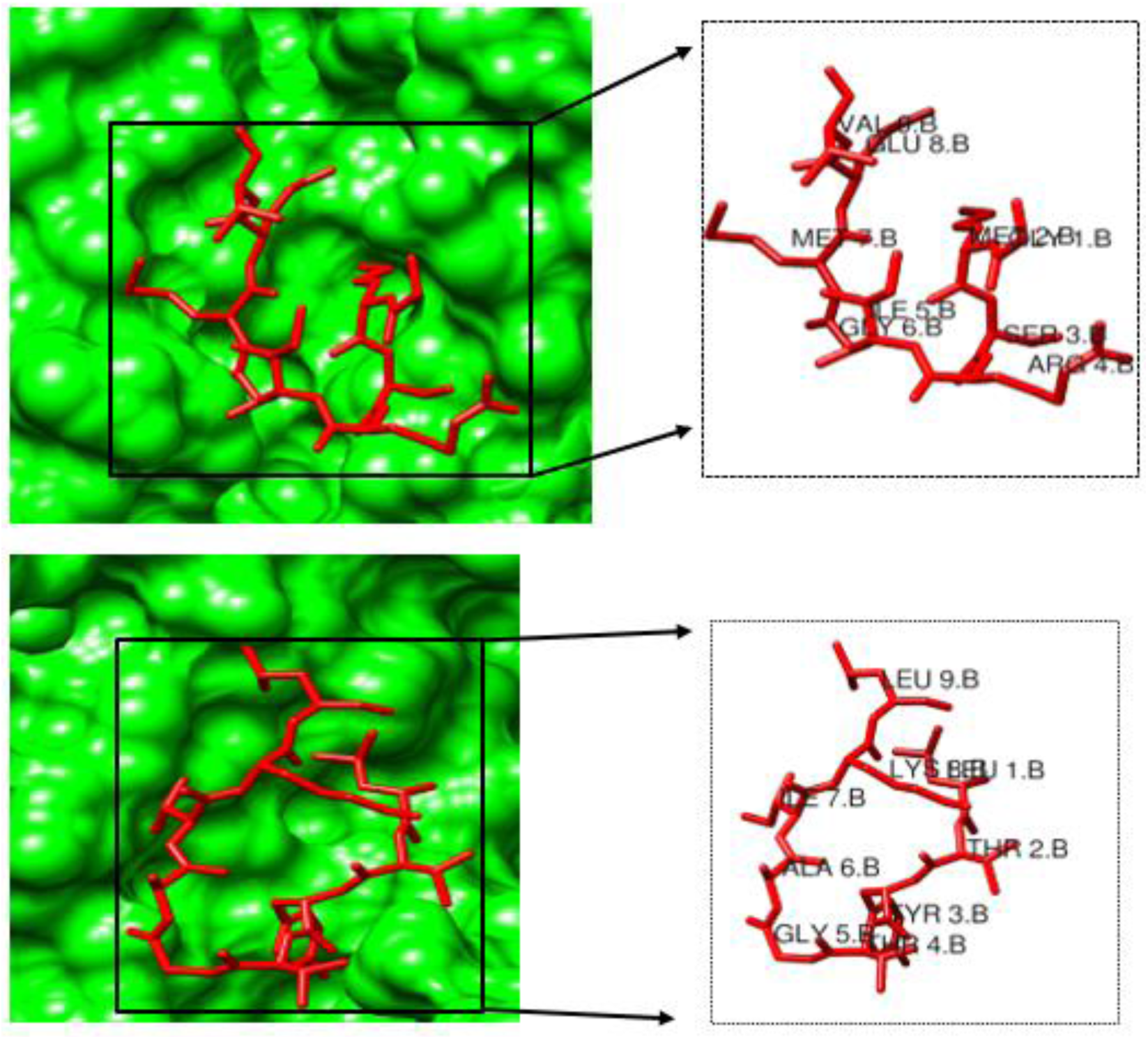
Molecular docking of the peptides “GMSRIGMEV and LTYTGAIKL, and the HLA A0201 molecule. GMSRIGMEV had binding free energy of −8.3 kcal/mol, and −10.0 kcal/mol for LTYTGAIKL.

### Eminent profiling of the chimeric vaccine construct

The final conjugated vaccine consists of two B cell linear epitopes, two CD4+, three CD8+ epitopes and a poly-histidine tag, which sums up the total of 78 amino residues. Given the high antigenicity index of 0.75 of the predicted vaccine, viable enough in eliciting both humoral and cellular immune responses, and their non-toxicity and allergenicity, an immuno-adjuvant to boost the antigenicity of the vaccine construct was excluded. The addition of poly-histidine residues at the C-terminal of the vaccine helps to convey increased purity to the recombinant protein and may contain specified epitopes that can be recognized by an antibody fragment, thereby increasing the efficacy and effectiveness of the peptide vaccine.

### Physiochemical and Solubility properties of the Vaccine

The molecular weight of the vaccine was 8558.93 Da and the bio-computed theoretical pI was 9.98, with an estimated half-life of 30 hours. The instability index was 30.09, signifying that the protein is stable (>40 signifies instability). The aliphatic index is computed to be 78.85, with a GRAVY score of −0.281, signifying its hydrophilic nature. The atomic composition of the vaccine is 376 carbon, 603 hydrogen, 115 nitrogen, 104 oxygen and 5 sulfur, thereby giving rise to the chemical formula C_376_H_603_N_115_O_104_S_5,_ with a total of 1203 atoms. The extinction coefficient at wavelength of 280nm was 2980 M^−1^ cm^−1^. The intrinsic vaccine solubility at a neutral pH 7 revealed the hydrophilic and hydrophobic core of the vaccine construct **[Figure 6]**.

**Figure 6:**
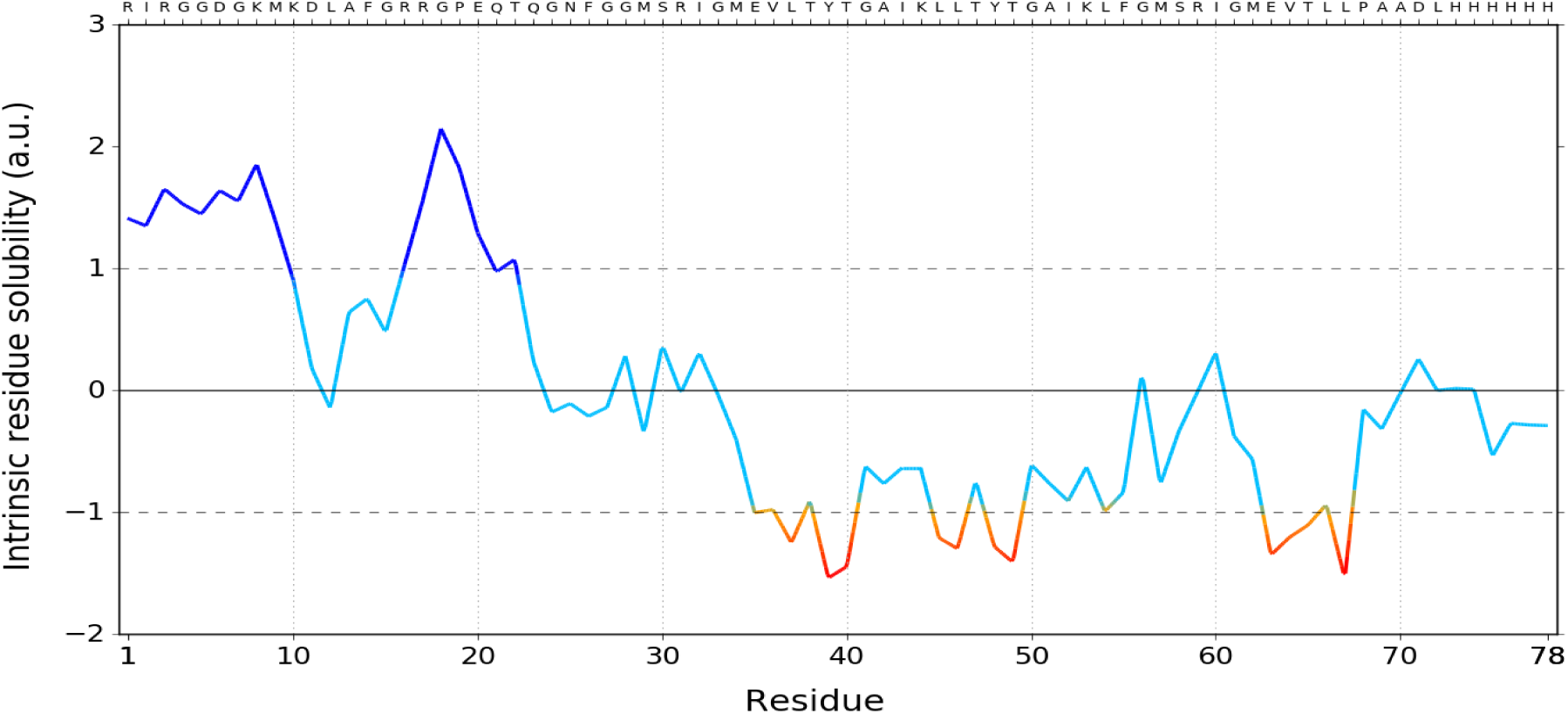
Solubility index profile of the peptide vaccine. Residues less than −1 depict the hydrophobic core of the vaccine peptide.

### Three-dimensional structure prediction of the Vaccine

The predicted model of the vaccine and its three-dimensional coordinate file was successfully obtained from I-TASSER [**Figure 7a**]. The results obtained from the server includes predicted secondary structure with a confidence score ranging from 0 to 9, predicted solvent accessibility, functional analogs protein, and binding site residues. The best model was selected with a C-score of −1.39, TM-score of 0.54±0.15, and RMSD at 6.3±3.8Å.

**Figure 7a:**
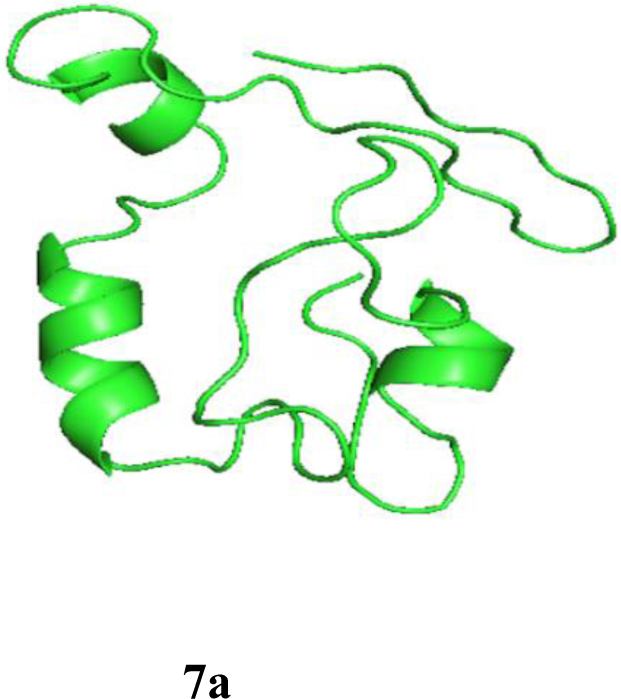
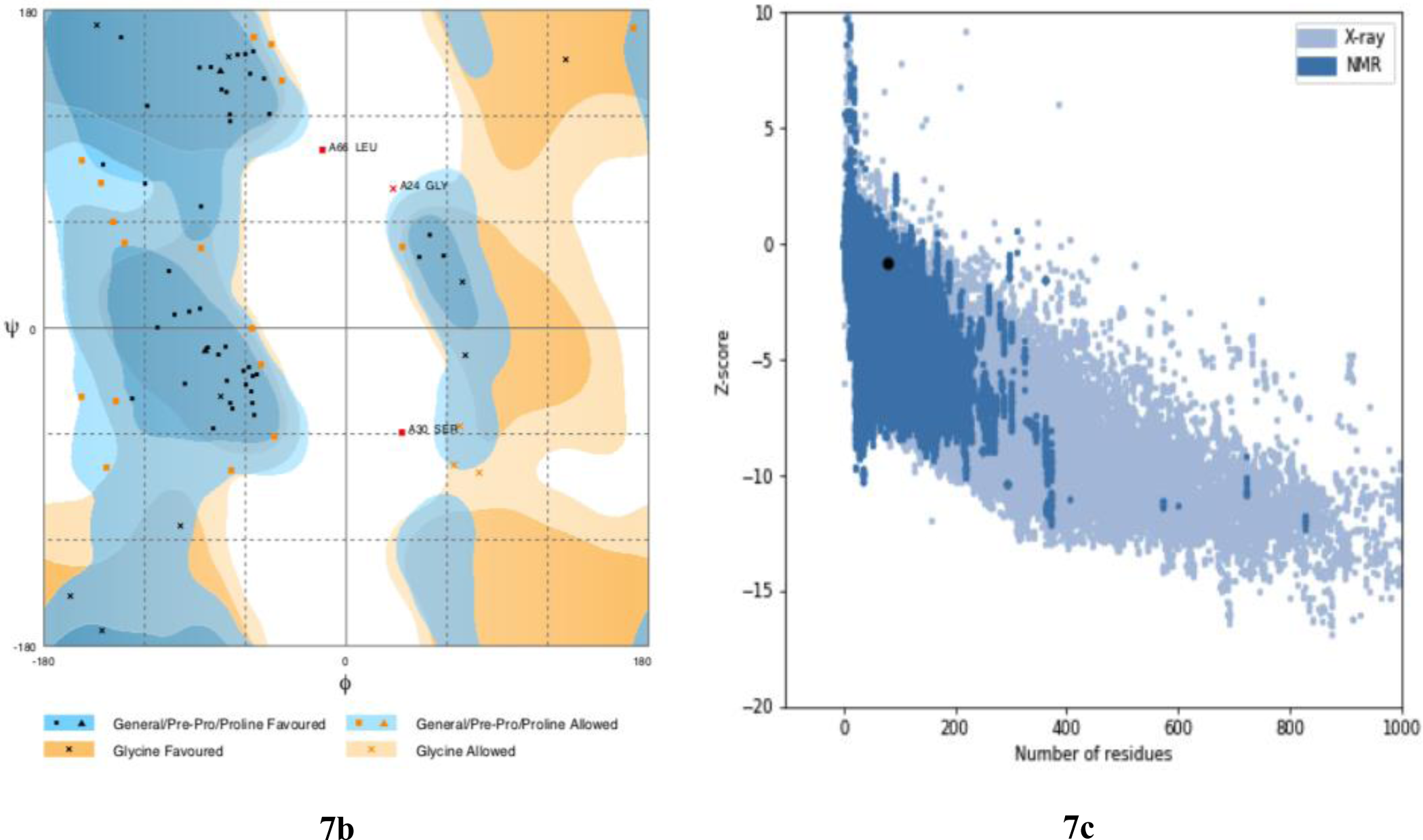
Three-dimensional structure of vaccine predicted by I-TASSER. Three-dimensional structure prediction validation of top score model of I-TASSER by **(7b)** RAMPAGE assessment of the Ramachandran Plot of selected model. Number of residues in favored region: 53 (69.7%), Number of residues in allowed region: 20 (26.3%), Number of residues in outlier region: 3 (3.9%), **(7c)** ProSA protein structure analysis results. Z score = – 0.79. Overall quality of the ultimate model is acceptable.

### Structural Validation of the predicted model

The Ramachandran plots of the predicted model were obtained to verify the stereochemical parameters of the protein structure. Ramachandran plot showed 69.7% residues in most favored regions and 26.3% residues in additional allowed regions i.e., the total of 96% residues in allowed regions which indicates a good quality model [**Figure 7b**], this was also attested by MolProbity Ramachandran plot which also showed 96.6% residues in allowed regions which also confirmed the quality of the predicted model [**Figure 7c**].

### Molecular interaction between the Peptide vaccine and the Toll-like receptor-5

A preliminary docking preparation was conducted by aligning the vaccine construct with the toll-like receptor to ascertain if both complexes are likely to interact. The TM alignment score obtained was 0.35240, which shows the likelihood of both proteins interacting with stable conformation. The docking structure of the human toll-like receptor-5 binding with peptide vaccine fragment was finally obtained. A conformational change occurs in the toll-like receptor-5 protein after binding with the antigenic peptide [**Figure 8**], with a binding energy of −8.6 Kcal/mol. The interface residue contacts characterizing both proteins interactions include: charged-charged was 7, charged-polar was 6, charged-apolar was 11, polar-polar was set at 0, polar-apolar was 7 and finally, apolar-apolar was 20 respectively. The non-interacting charged surface was 20.26%, while the non-interacting apolar surface was 42.41%. The interacting amino residues of the vaccine comprising of ASN 25, PHE 26, GLY 27, GLY 28, VAL 35, LEU 36, and THR 37, was able to form hydrogen bond with PRO 604.A, ASP 607.A, CYS 646.A, LEU 650.A, PHE 653.A, LEU 654.A, LEU 642.B, CYS 646.B, and THR 649.B of the receptor.

**Figure 8:**
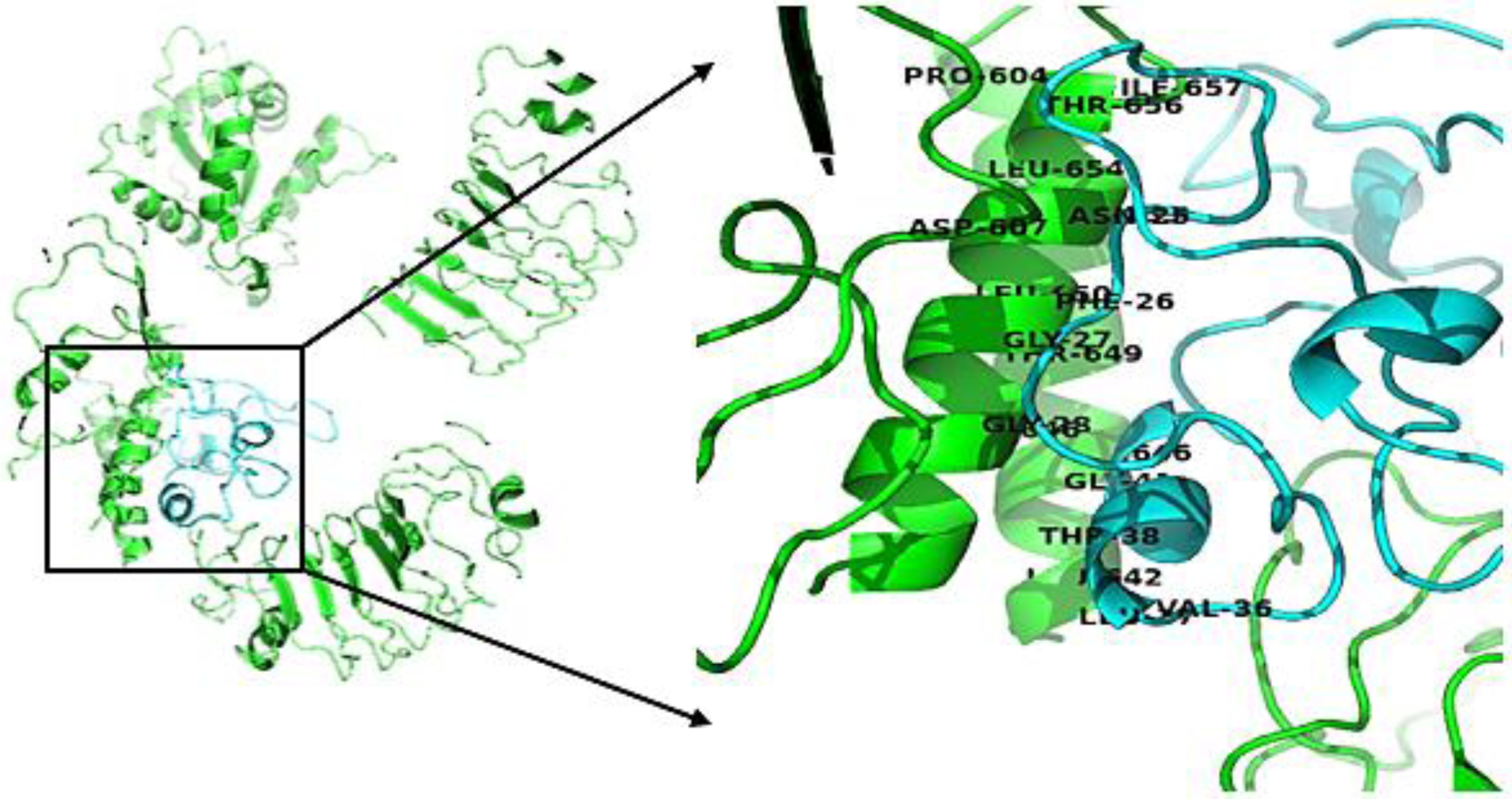
The interaction of the proposed vaccine construct with TLR5. The vaccine and TLR5 are shown in red and blue color respectively. The blue color signifies the vaccine, while the green color also signifies the chain of the receptor.

### Molecular dynamics simulations

The interaction between the peptide vaccine and the toll-like receptor was scrutinized to check for their protein stability and deformation. This analysis relies on the associated coordinates of the docked protein complex **[Figure 9a-9d]**. The eigen value found for the complex was 2.871961e-06. The low eigen value for the complex signifies easier deformation of the complex, indicating that the docking analysis between the vaccine and the TLR5 will activate immune cascades for destroying the viral antigens.

**Figure 9a-9d.**
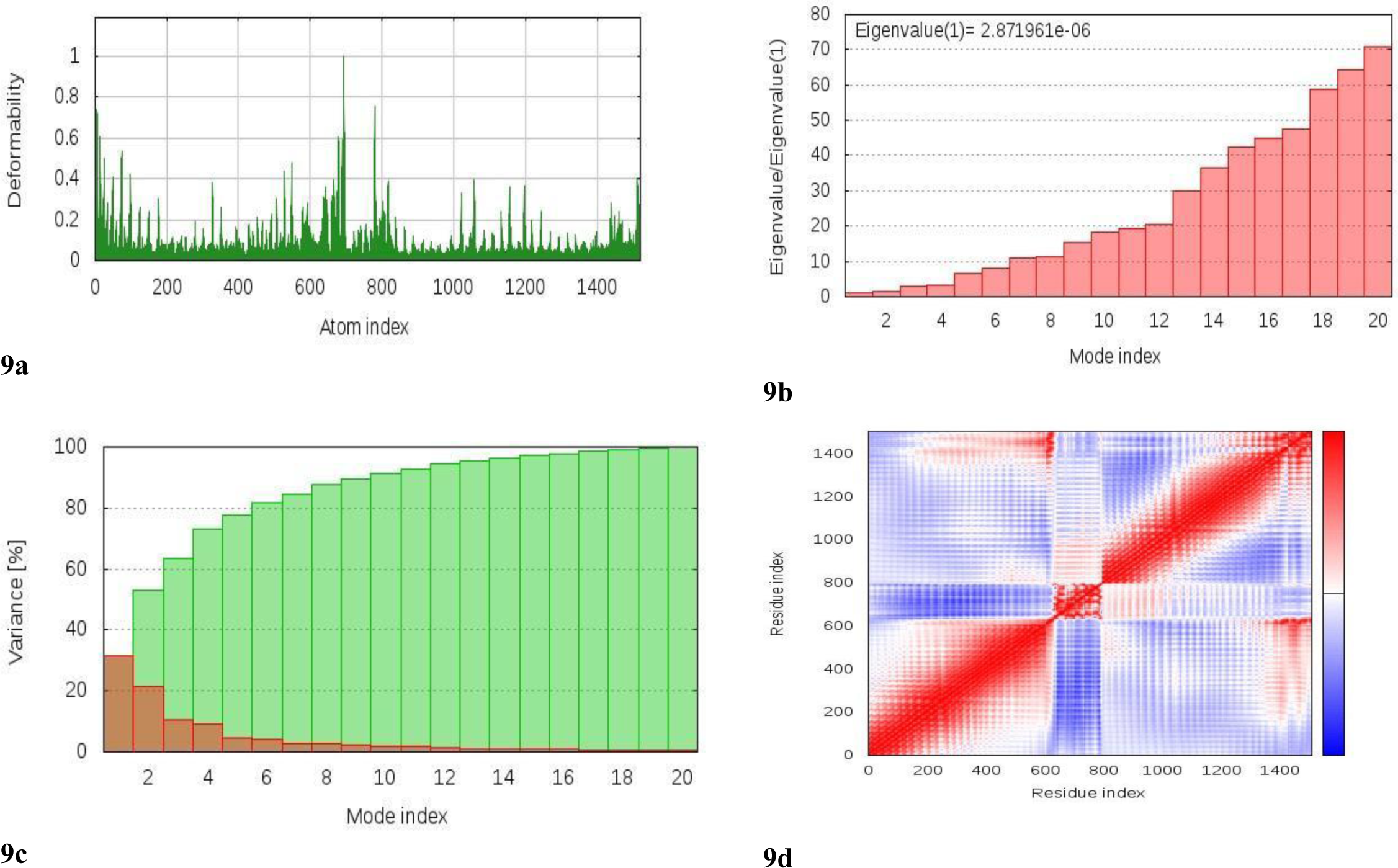
Molecular dynamics simulation of the vaccine-TLR5 complex, showing (a) eigen value; (b) deformability; (c) B-factor; (d) Covariance matrix indicates coupling between pairs of residues, i.e. whether they experience correlated (red), uncorrelated (white) or anti-correlated (blue) motions.

### Codon Optimization and *In Silico* Cloning

The length of the optimized vaccine codon sequence was 234 nucloetides. The GC content of the improved cDNA sequence was 54.27%, which still falls within the recommended range of 30-70%, for effective translational efficiency. The codon adaptive index was calculated as 1.0, falling within the range of 0.8-1.0, signifying the effective expression of the vaccine construct in the *E. coli*. EagI-NotI and SAlI sites were subsequently cloned into the pET28a D315A vector. The estimated length of the clone was 6.954 kbp [**Figure 10**].

**Figure 10:**
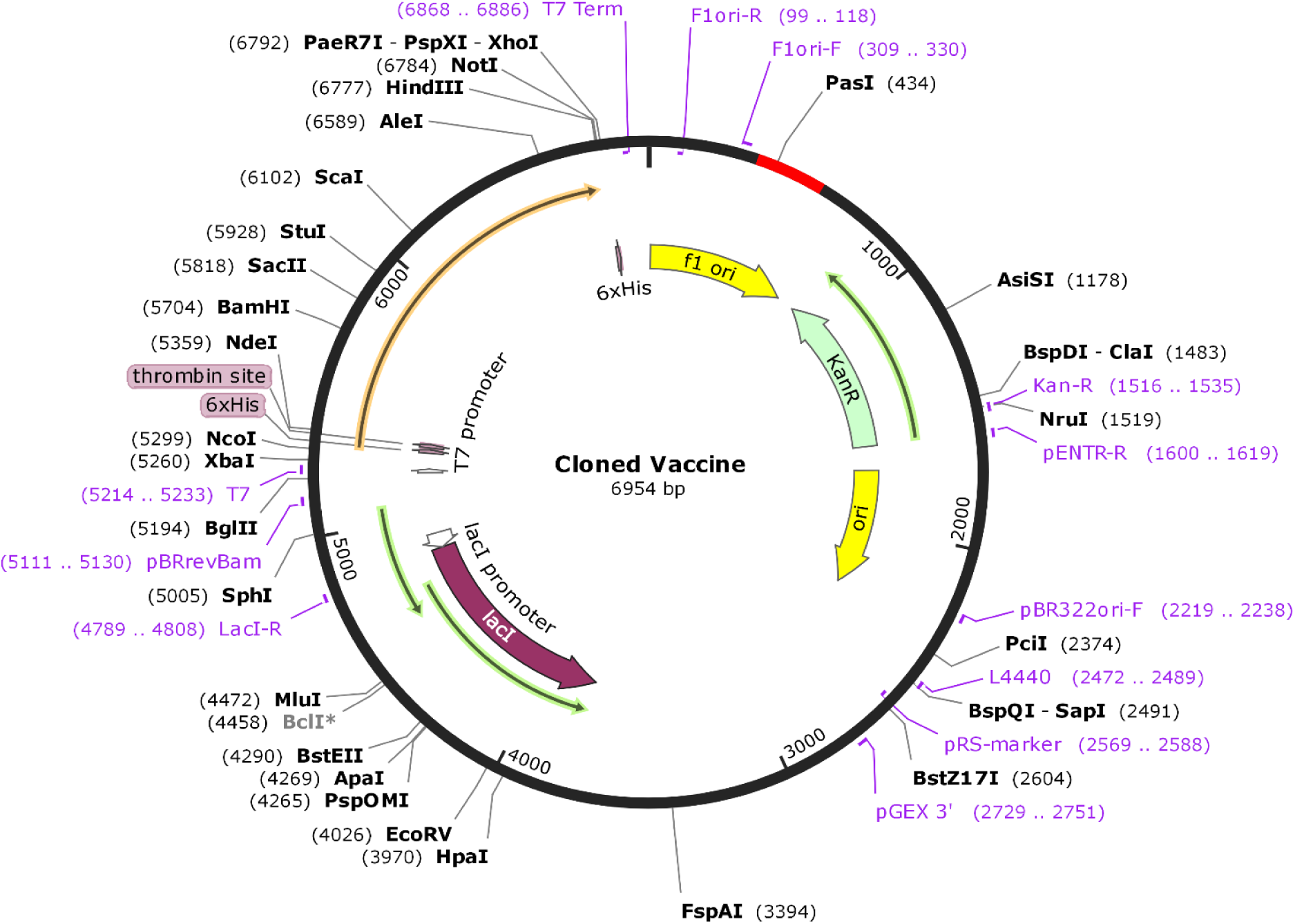
*In silico* cloning of the final vaccine construct into expression vector where the red part indicates the coding gene for the vaccine enzymatically restricted by PasI.

## Discussion

The menace of the coronavirus pandemic on global health has made imperative the development of safe, stable, and effective vaccines against it. Currently, most vaccine designs against the SARS-COV-2 virus are targeted against the Spike protein either as whole inactivated or attenuated viruses because of its ability to elicit neutralizing antibodies to block virus-receptor interaction and neutralize viral infection of cells [**25, 26**]. However, targeting the S protein may induce high neutralizing antibodies which are incapable of inducing long-lasting protection against the virus. Hence, alternative viral proteins may be considered. The N protein does not elicit neutralizing antibodies but may induce specific antibody and cellular immune responses [**27, 28**]. Also, the N protein is more conserved and induces long-lived memory T-cells in humans, features that make it a potent vaccine candidate [**29-31, 28**]. In this study, we explored the potentials of the Nucleocapsid phosphoprotein RNA binding domain as a subcomponent vaccine candidate which could induce protective immunity against the SARS-COV-2 virus.

Structural alignments show that the N protein is less conserved. This could be attributed to the mutation of the N protein sequence in the coronavirus family. Dawood (2020), reported a moderate mutation of the N-protein of coronaviruses [**32**]. This is indicative that the SARS-COV-2 virus may have originated from mutations within the family as has been previously suggested [**33**].

The humoral immune response plays a clear role in vaccine-mediated defense against infections given their role in maintaining memory cells, prolonged survival, and sentinel against re-infection [**34**]. Subcomponent vaccines that are capable of focusing the humoral immune response on specific antigenic epitopes can be predicted to be crucial and most beneficial in inducing specific antibodies and long-lived memory immune cells. Here, we predicted 2 viable antigenic regions using diverse investigation tools and processes for the calculation of the B-cell linear epitopes. These 2 regions (one a dodecapeptide and the other, a pentadecapeptide) were selected based on their conservancy, high antigenicity, non-toxicity, and non-allergenic properties. Due to findings from the diverse investigation tools employed, we report that our identified epitopes can elicit humoral immunity. This is in consonance with previous studies [**35, 36**] who reported the ability of the SARS-COV N protein to elicit N-specific humoral immune response *in vitro*. Also, the discontinuous B-cell epitope prediction, which are amino acid residues that were brought into close proximity within the folded protein structure [**37**] revealed that the chain D component of the protein had the highest propensity score. This is important because antibody binding is not just determined by the linear peptide segment but is also influenced by adjacent surface regions [**38, 39**].

The outcome of disease infection in humans is usually a factor of the strength of the immune response mounted against the infectious agent and this is usually orchestrated by MHC molecules of the cellular immune system [**33**]. While B cells recognize epitopes on the surface of the infectious agent, T cells recognize epitopes on MHCs. Two subpopulations of T cells (Cytotoxic T lymphocytes and Helper T lymphocytes) are involved in epitope recognition and while CD8+ CTLs recognize antigens presented on MHC class I, CD4+ HTLs recognize antigens on MHC class II. Also, CD4+ HTLs play a vital role in coordinating both humoral and cell-mediated immune responses [**40-42**]. The two CTLs (C_2_ and C_3_) were selected due to their high antigenicity among the four identified epitopes. The three HTL epitopes (H_2_, H_4_ and H_6_) were selected based on their non-allergenicity and high antigenicity. Even though other epitopes had higher antigenicity scores, they were however found to be allergenic, able to elicit a harmful autoimmune response. Therefore, they were not selected.

Interestingly, a recent study predicted some T cell epitopes which can be recognized by MHC class II CD4+ HTL alleles of the Asian and Asian-pacific region population [**33**]. While they considered only the MHC class II epitopes, we considered both MHC class I and II epitopes in our study. Also, no HTL epitope were identified in their Nucleocapsid protein, however, we report that two CTL and three HTL epitopes with high antigenicity and absence of allergenicity were identified on the Nucleocapsid protein N terminal RNA binding domain of SARS-COV-2. Also similar to our study, Wang *et al*. (2003) predicted 4 strong antigenic sites and synthesized 2 strong immunogenic peptides on the N-protein of SARS-CoV of which, one of our predicted peptides falls within the synthesized peptides showing medium-strong immunogenicity [**43**]. However, our study also revealed epitopes which have not been reported earlier. This indicates that the N protein is one of the major antigens of the coronaviruses with diverse potent epitopes for vaccine development.

We examined the interaction of the designed vaccine with TLR5 using molecular docking analysis. The binding interfaces between the TLR5 and the peptide vaccine consisted of hydrogen bonding and hydrophobic interactions. Also, the relative binding free energies of the vaccine-TLR5 complex suggests that the linking of the peptide vaccine construct to the receptor elicits conformational changes that favor stimulation of the TLR5 immune molecules and indicate a favorable protein-protein interaction of our vaccine construct with the innate immune receptor. Similarly, the result of the molecular simulation analysis to examine for stability and deformation in the interaction between the vaccine and the TLR5 complex found a low eigen value for the complex indicating easier deformation of the complex. This implies that the docking analysis between the vaccine and the TLR5 will activate immune cascades for destroying the viral antigens. Even though further *in vitro* and *in vivo* analysis is proposed to corroborate this result.

We therefore report that the *in silico* epitope-based vaccine construct targeting the SARS-COV-2 N protein N-terminal RNA binding domain shows prospects as a potent, safe, and effective candidate with high antigenic properties and a balanced immune response operating through both innate and adaptive pathways.

## Conclusion

The lack of an effective therapeutic candidate against the novel coronavirus has created a huge task for biomedical researchers to seek research approaches for overcoming the pandemic. This study was designed in furtherance of steps towards vaccine development. We used the primary amino acid sequence of the SARS-COV-2 to design a subcomponent peptide vaccine construct. The vaccine construct has both adaptive (B and T cell) epitopes as well as a favorable interaction with the pathogen recognition receptors (PRRs) (via the TLR5) of the innate immune system. Each of the predicted epitopes has antigenic properties in the absence of allergenic properties. Generally, this study applied a series of immunoinformatic tools to predict a safe, stable and effective peptide vaccine that may fight against the SARS-COV-2 viral infection. However, we propose experimental validations to prove this computational work.

## Acknowledgment

None

## Supplementary Data

| S/N | PDB ID | Protein name | Year Released |
| --- | --- | --- | --- |
| 1. | 6M3M | Crystal structure of SARS-CoV-2 nucleocapsid protein N-terminal RNA binding domain | 2020 |
| 2. | 6WKP | Crystal structure of RNA-binding domain of nucleocapsid phosphoprotein from SARS CoV-2, monoclinic crystal form | 2020 |
| 3. | 6YI3 | The N-terminal RNA-binding domain of the SARS-CoV-2 nucleocapsid phosphoprotein | 2020 |
| 4. | 6VYO | Crystal structure of RNA binding domain of nucleocapsid phosphoprotein from SARS coronavirus 2 | 2020 |
| 5. | 1SSK | Structure of the N-terminal RNA-binding Domain of the SARS CoV Nucleocapsid Protein | 2004 |
| 6. | 2OFZ | Ultrahigh Resolution Crystal Structure of RNA Binding Domain of SARS Nucleopcapsid (N Protein) at 1.1 Angstrom Resolution in Monoclinic Form. | 2007 |
| 7. | 4UD1 | Structure of the N Terminal domain of the MERS CoV nucleocapsid | 2015 |
| 8. | 6KL2 | Structure of the N-terminal domain of Middle East respiratory syndrome coronavirus nucleocapsid protein | 2020 |
| 9. | 3HD4 | MHV Nucleocapsid Protein NTD | 2009 |
| 10. | 4LM9 | Crystal structure of HCoV-OC43 N-NTD complexed with GMP | 2014 |
| 11. | 4J3K | Structure of the N-terminal domian of human coronavirus OC43 nucleocapsid protein | 2013 |
| 12. | 2GEC | Structure of the N-terminal domain of avian infectious bronchitis virus nucleocapsid protein in a novel dimeric arrangement | 2006 |
| 13. | 2BXX | Crystal structure of the N-terminal domain of IBV coronavirus nucleocapsid. Native crystal form | 2005 |
| 14. | 2BTL | Crystal structure of the N-terminal domain of IBV coronavirus nucleocapsid | 2005 |
| 15. | 5N4K | N-terminal domain of a human Coronavirus NL63 nucleocapsid protein | 2017 |

